# Palaeogenomics reveals 1,500 years of population history of the peoples of the Chonos Archipelago, Chile

**DOI:** 10.1101/2025.11.26.690513

**Authors:** Roberta Davidson, Omar Reyes, Xavier Roca-Rada, Epifanía Arango-Isaza, Kalina Kassadjikova, Chiara Barbieri, Bastien Llamas, Lars Fehren-Schmitz

## Abstract

The Chonos archipelago, located in northwestern Chilean Patagonia, was historically inhabited by an Indigenous sea-faring population known to the European colonists as the "Chono". Previous research has contextualized the human occupation of this region with radiocarbon dates and ancient mitochondrial genomes, offering a partial perspective on the history of its inhabitants. Here we present a paleogenomic analysis of 20 ancient human individuals from 6 archaeological sites, dated between 1,600-50 years before present (BP). We successfully captured over 100,000 SNPs from 15 of 20 individuals and recovered 12 full mitogenomes. All individuals presented unadmixed Indigenous South American ancestry and formed a separated genetic cluster relative to other ancient and modern South American genomes, indicating a unique Chonos ancestry. This Chonos ancestry falls within a broader cluster of late-Holocene Patagonian ancestry, most similar to the Kawésqar peoples who neighbour to the south. Within the Chonos, we distinguish a northern and southern ancestry cluster. The northern Chonos cluster exhibits some genetic connections to the present-time inhabitants of the neighbouring island of Chiloé, who are connected to Huilliche Indigenous history. Our findings reveal a point of contact between southern Chonos/Patagonian ancestry to the south and Mapuche ancestry to the north, and confirm that sea-faring subsistence was a knowledge transferred between Patagonian peoples.

## Introduction

The peopling of the Americas was one of the last human continental migrations. Ancient and modern genetic data places this early expansion between 23 kya and 16kya ^1^ predominantly along a coastal route starting from Beringia and following along the Pacific Coast of the Americas ^2^. Humans rapidly reached the Southern Cone of South America (SCSA)—the large cone-shaped geographical region encompassing Chile, Argentina, Uruguay and southern parts of Paraguay and Brazil (Figure 1A). Revised radiocarbon dating of archaeological evidence at Monte Verde, Chile, dates first human occupation in the SCSA to 18.5–14.5 kya ^3^, although the most robust estimates are closer to 14.5 kya ^4^. Concurrently, comprehensive analysis of radiocarbon dates suggests a contemporaneous peopling of Chile approximately 13,000 cal BP (calibrated years before present) ^5^.

**Figure 1:**
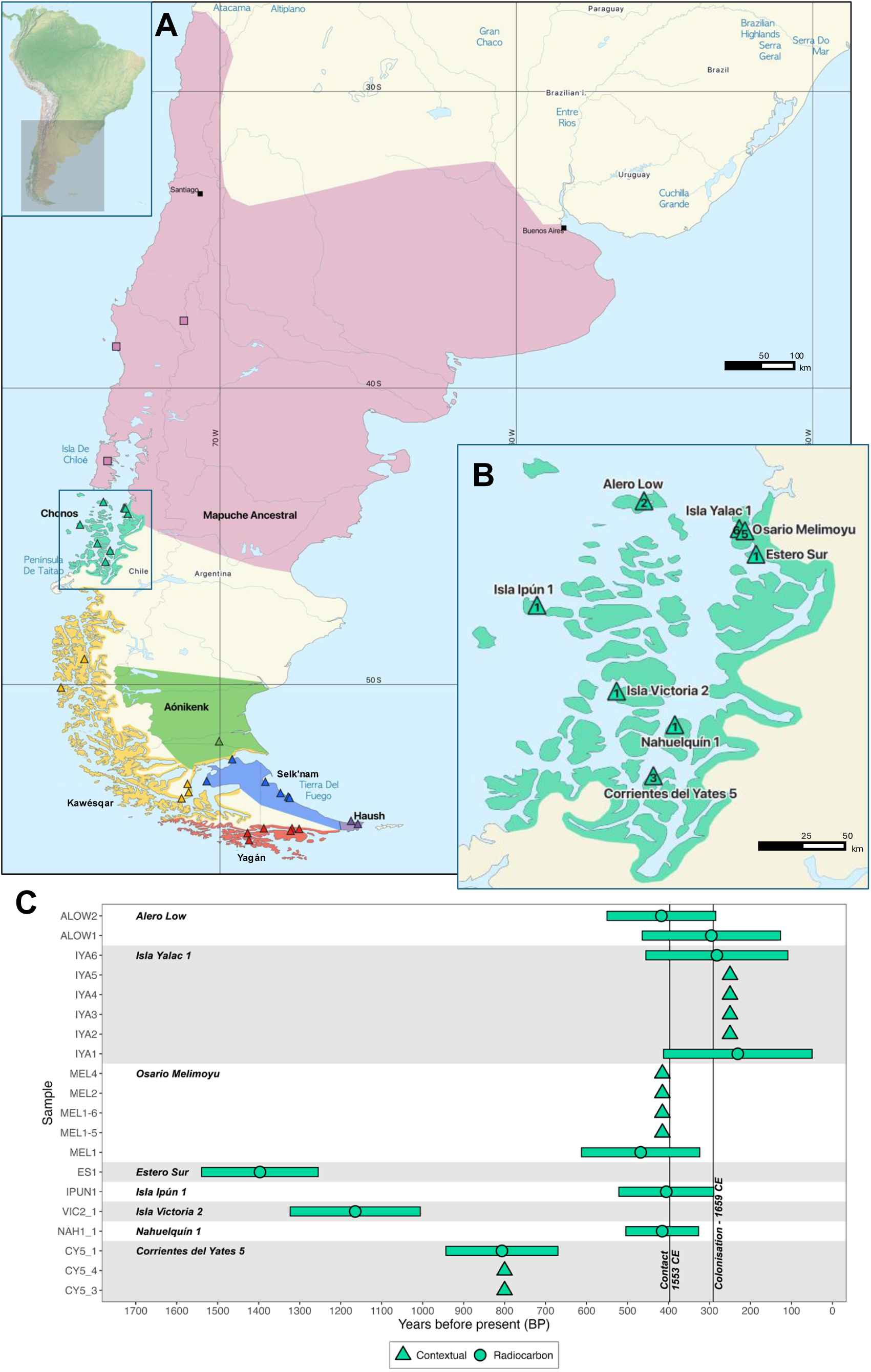
(A) Map of the Southern Cone of South America, with ancestral population regions (highlighted with colours) and sampling locations used in this study (squares: present-day populations; triangles: archaeological sites). Linguistic-cultural regions redrawn from ^12,17^. (B) Location of study sites within the Chonos archipelago labelled with the number of individuals from each site (triangles). (C) Chronology of novel individuals sampled from the Chonos archipelago analysed in this study, grouped by archaeological site and ordered by latitude. Calibrated radiocarbon dates are presented with error bars, contextual dates as points (triangles). Black vertical line indicates the dates of earliest European contact in Patagonia (1553 CE) ^34^ and colonisation of Patagonia (1659 CE).

Within the SCSA, Patagonia is the southernmost region, characterised by diverse archipelagos of islands along the south-western coast of the continent and the southern plains of Argentina. Most inhabitants of south-western Patagonia adapted to sea-faring ways of life, as evidenced by maritime technology such as canoes that begin to appear in the archaeological record by 7.3 kya in the southernmost region of Tierra del Fuego ^6,7^. When Europeans invaded southern Patagonia in the 16th century, they reported three major Indigenous populations of maritime peoples; Yagán, Kawésqar, and Chono (Figure 1A) ^8–11^. Previous research based on mtDNA and genome-wide variation suggests a shared origin for the southernmost late maritime populations, as well as long-term continuity for at least 2 kya ^12,13^. Seafaring subsistence has been recorded for the whole Pacific coastal portion of the Southern Cone, up to the Chonos archipelago, where studies of dietary isotopes in pre-contact Chono individuals show signals of a seafood diet for, confirming archaeological observation of seafood consumption ^14–16^. Recent palaeogenomic research indicates that sea-faring technology and culture adaptations in southern Patagonia were independently regionally developed rather than introduced via immigration ^17,18^, and that genetic continuity has persisted since ∼6.6 kya with some later gene flows occurring prior to 2 kya ^17^

The people referred to in ethnohistory as “Chono” inhabited the Chonos archipelago in northwest Chilean Patagonia ^8,9,19,20^, although ethnohistorical sources indicate a great diversity of Indigenous peoples in the region ^21,22^. The Chonos archipelago (43°–45°S) is a collection of over 150 fjordic islands between the island of Chiloé to the north (41°–43° S) and the Taitao Peninsula to the south (45°S–47°S) (Figure 1A) ^14^. The archipelago receives over 3,000 mm of rain per year ^23^ and has an average daily temperature of 10°C ^24^. The region is subject to cyclic glaciation and volcanic activity that has shifted coastlines throughout the Holocene and hence impacted preservation of archaeological sites ^14,24^. Recently, the first human ancient DNA study from the Chono individuals reported ancient mitogenomes characteristic of the SCSA (predominately haplogroups D1 and C1), and most similar to those found in ancient Kawésqar peoples immediately to the south ^18^.

To the north of the Chonos archipelago, neighbouring populations include the Mapuche/Huilliche people, who today are one of the most represented Indigenous groups in the SCSA, maintaining a significant presence in southern Chile and parts of Argentina. Historically, their ancestral range extended from Chiloé Island to central Chile and reached into Argentina (Figure 1A). The present-day Mapuche self-identify according to the region they inhabit, for example as Lafkenche along the Pacific coast, Pehuenche in the Andean mountains, and Huilliche in the southern regions and Chiloé island, where some communities remark a distinction between Huilliche and Mapuche history, language and cultural identity ^25^. Recent studies of genome-wide data from living populations of Mapuche/Huilliche descent link their ancestry to local aDNA from the mid-Holocene (∼5kya) and suggest a broad genetic relationship with other ancient and modern SCSA populations^12,25^.

Following European colonisation in the 16th century, drastic population declines are recorded throughout the Americas ^26^. In particular, a 96% population loss was estimated among the Huilliche–Pehuenche communities in Southern Chile ^27–30^. While some people living today do identify as Yagán and Kawésqar, the impact of European occupation was completely devastating for the Chono ^10,31^. Within just two centuries, they experienced a severe population decline, with many individuals being enslaved and forcibly relocated, as far away as central Chile and even Peru ^14^. This resulted in the depopulation of the Chonos archipelago and the extinction and loss of Chono culture ^32,33^, leaving the Chono as one of the least understood peoples to have inhabited Patagonia ^14,18,24^.

The lack of genome-wide data from the ancient Chono leaves unresolved whether maritime technologies in this region spread through cultural exchange or migration from neighbouring Patagonian groups. Furthermore, potential connections between the Chono and Mapuche peoples, particularly the Huilliche from the neighbouring island of Chiloé, remain unknown. Answering these questions is key to understanding dynamics between ancient populations in the SCSA, at the interface between maritime and terrestrial cultures. Here we report genome-wide data from 20 individuals from 8 sites in the Chonos archipelago (Figure 1B), spanning 1600 years (Figure 1C, Table S1). We characterise the Chono population from palaeogenomic evidence and investigate the relationships between the Chono and other populations in the SCSA.

## Methods

### Samples

We generated genome wide data for 20 ancient individuals from 8 different archaeological sites in the Chonos archipelago, Chile (Figure 1B). Archaeological sites are described in detail by Reyes ^14^. The individuals span approximately 1600 – 200 cal BP (Figure 1C) according to radiocarbon dating and contextual dating conducted previously ^14^. However, these radiocarbon dates have since been recalibrated, potentially placing the earliest individuals into post-European-contact time (Figure 1C). Given that we do not identify any non-American genetic ancestry component we cannot genetically establish if the individuals actually experienced European contact, but we acknowledge that the lives of the studied individuals could be impacted by these events. Nevertheless, we do not interpret the demographic history recorded through the lens of colonisation.

### Ethics and community engagement

The ancient individuals analysed in this chapter were made available to us, in accordance with the requisite local and institutional permissions, through a collaboration with Chilean archaeologist and present co-author, Omar Reyes, whose extensive career has been dedicated to research in the Chonos Archipelago ^14^. It is important to note that there is no known contemporary community claiming descent from the ancient Chono peoples ^32,33^. However, this does not preclude the possibility of Chono genetic ancestry persisting within living populations that have lost connection to this culture and identity, nor does it reduce the importance of this research to the contemporary Indigenous peoples of Chile.

The project builds on previous community engagement work in southern Chile during a genetic study of Mapuche history ^25^. One of the questions put forward by the participants and community members was about a possible ancestral connection between the island of Chiloé and the Chono neighbours, who might have left linguistic traces (in toponyms, surnames and local terminology) as well as cultural traces in Chiloé ^35^. We prepared a report in Spanish to be shared with local communities to disseminate our results (Supplementary Document 2) With this we demonstrate one approach to transparent production and dissemination of scientific knowledge and community engagement, acknowledging the complexity of regional cultural context and the critical perspectives of Indigenous people of the Southern Cone.

### DNA extraction and sequencing library preparation

DNA was extracted from tooth root powder using a method optimised to retrieve highly degraded ancient DNA fragments ^36^ but with a modification to the protocol that includes a bleach pre-wash step ^36,37^. From each DNA extract, we prepared Partially UDG-treated ^38^ single-stranded DNA libraries ^39^. These steps were performed at the dedicated ancient DNA facilities of University of California, Santa Cruz (UCSC) Human Paleogenomics Lab.

### Library enrichment and sequencing

Libraries were enriched using two different assays. First, the Prime Plus enrichment kit (Daicel Arbor Biosciences), which includes DNA baits targeting ∼1.2 million nuclear SNPs and the whole mitochondrial genome. However, after discovering the allelic bias of this assay ^40^, we repeated enrichment with the Twist Biosciences enrichment kit ^41^, which includes DNA baits targeting the same ∼1.2 million nuclear SNPs as well as additional SNPs and tiling regions ^42^. In the final analyses, uniparental marker sequences (mitochondrial and Y chromosomes), captured with the Prime Plus are used as they are not impacted by the allelic bias discovered in the Prime Plus enrichment kit ^40^. For genome-wide analyses, only the 1.2 million nuclear SNPs captured with the Twist reagent are used, as they are less impacted by allelic bias than the Prime Plus, and exhibit approximately similar levels of bias to the widely-accepted 1240k enrichment assay ^41^.

To prepare for enrichment with the Prime Plus kit, libraries were over-amplified in order to reach the minimum required DNA input of 500 ng per sample. Libraries were then grouped into equimolar pools of 2-4 samples based on similar endogenous DNA %. A total of 2000 ng of pooled libraries in 10µl was used in each capture reaction. Enrichments were performed in accordance with the manufacturer’s protocol at the Australian Centre for Ancient DNA, Adelaide. Libraries enriched with the Prime Plus reagent were paired-end sequenced on an Illumina NovaSeq 6000 (2x100bp), performed by the Kinghorn Centre for Clinical Genomics (Sydney, Australia).

To prepare for enrichment with the Twist Biosciences kit, libraries were similarly over-amplified and then grouped into equimolar pools of 2-4 samples based on similar endogenous DNA fraction. A total of 1000 ng of pooled libraries was dehydrated and input to each capture reaction. Library pools were enriched according to the manufacturer’s protocol at the UCSC Human Paleogenomics lab. Libraries enriched using the Twist reagent were paired-end sequenced (2x100bp) on a NovaSeq X (Illumina) at Fulgent Genomics (Los Angeles, USA).

### Data processing

All ancient DNA sequencing data were demultiplexed with bcl2fastq 2.19.1 and processed using the nf-core/eager pipeline v2.4.5 ^43^. Reads with a length of 25 nucleotides or more were mapped to the GRCh37+decoy reference genome (hs37d5) using bwa-aln v0.7.17-r1188 ^44^ with ancient DNA parameters ^45^ and mapping quality filter 30. Genetic sex was called with Sex.DetERRmine v1.1.2 ^46^. Pseudohaploid genotypes were called at the enrichment target sites using the random haploid mode in pileupCaller from sequenceTools v1.5.2 ^47^. Ancient DNA damage was assessed using DamageProfiler to assess read length distributions and quantify the proportion of cytosine to thymine misincorporations at the 5’ end of sequenced DNA molecules and the G to A misincorporations at the 3’ end.

### Uniparental markers

Y chromosome haplogroups were called for male individuals using YHaplo ^48^. Mitochondrial genomes were mapped to the mitochondrial Cambridge reference sequence ^49^ using nf-core/eager ^43^. Haplocheck 1.3.3 ^50^ was used to report mitochondrial contamination levels and assign haplogroups. Read pileups were then manually inspected in Geneious (11.1.5) and polymorphisms were called at positions with ≥ 3X coverage and variant frequency ≥ 0.7. Consensus FASTA sequences were generated in Geneious Prime with a threshold of 50% and N called for coverage < 3X. Haplogroups were then assigned again with Haplogrep 3 ^51^ on PhyloTree 17 - Forensic Update 1.2 ^52,53^, and a mitochondrial genome graph method, haploCart ^54^. Genome sequences with discrepancies between Haplogrep 3 and haploCart assignment methods were manually inspected in Geneious for haplogroup diagnostic SNPs and assigned to the appropriate haplogroup.

### Genetic relatedness

To explore close genetic relatedness between individuals, we used BREADR, a pairwise mismatch rate (PMR) based kinship detection method ^55^ was run on pairwise combinations of Chono individuals that had ≥ 100,000 SNPs. A filter length of 1e4 was used to calculate counts. Relationships up to the 10th degree were tested using the *test_degree* function for pairs from the same archaeological site and time period.

### Comparative datasets

Newly sequenced ancient samples were analysed together with comparative genomic datasets, which depending on the analysis include ancient and modern genetic profiles from the Americas, and with population references from other continents ^1,12,17,25,29,56–75^. Full details of each comparative dataset are detailed in Table S2.

### Runs of homozygosity

To investigate runs of homozygosity, the package hapROH ^76^—designed to infer ROH blocks from pseudohaploid ancient genomes—was run on ancient individuals from the SCSA. Samples were filtered to have at least 400,000 SNPs covered in the 1240k target set, identified using *--missing* in Plink 1.9 ^77,78^. The 1000 Genomes data were used as the reference dataset with the model parameter *e_model=“haploid”*.

### Conditional heterozygosity

A conditional heterozygosity analysis was performed on all ancient SCSA groups with n > 1 population size using the Popstats package ^72^ September 27, 2018 version, with the *--pi* flag and otherwise default settings.

### Population structure

Genotype data were first pruned for linkage disequilibrium using the Plink 1.9 flag *--indep-pairwise 200 25 0.4*, indicating a window size of 200 variants, step size of 25 variants and an *r^2^* threshold of 0.4, which resulting in a dataset of 170,094 SNPs. Principal components analysis (PCA) was computed using EMU ^79^ software, run with a minor allele frequency filter of 0.01. The explanatory percentage of each eigenvector was calculated as the eigenvalue out of the sum of all eigenvalues outputted. Population structure was then assessed with ADMIXTURE 1.3 ^80^, run unsupervised from K=2 to K=12 with 10 replicates each. Results were visualised using PONG v 1.5 ^81^ with a similarity threshold of 0.9 between replicates.

### Outgroup ***f*_3_** statistics and population clustering

All *f*_3_ statistics were computed using ADMIXTOOLS 7.0.2 ^82^ qp3Pop with the parameter *inbreed: YES,* to account for pseudohaploid genotypes. Population *f*_3_ statistics were computed in the configuration *f*_3_(Mbuti; Chono, X) to measure each population’s shared drift with the Chono population. Population clustering was performed by computing *f*_3_(Mbuti; Ind1, Ind2) pairwise for all individuals. A Multidimensional Scaling Plot (MDS) was then drawn from a matrix of the statistic 1-*f*_3_(Mbuti; Ind1, Ind2) using the *cmdscale* function with *k=10* parameter from the R stats package ^83^. Similarly, clustering heatmaps of pairwise relatedness were drawn from the matrix of pairwise *f*_3_ statistics using the pheatmap package^84^.

### *f*_4_ statistics

All *f*_4_ statistics were computed using ADMIXTOOLS 7.0.2 ^82^ qpDstat with *f4mode: "YES* and the parameter *inbreed: YES* to account for pseudohaploid genotypes.

### qpAdm

We used the qpAdm program from the ADMIXTOOLS v5.1 package ^82^, with the *allsnps: YES* option to minimally reduce the number of SNPs used and subsequently increase the power to reject models, to model the ancestry in our newly reported individuals. Hence, we quantified the proportion of genetic ancestry contributed by each source. The ancestry proportions in the target population are inferred on the basis of how the target population is differentially related to a set of reference/outgroups via the source populations. For all the models applied here, we have used a set of 6 outgroups (Mbuti.DG, CHB.DG, China_Tianyuan, Russia_Ust_Ishim_HG.DG, USA_Ancient_Beringian.SG, USA_Anzick_realigned.SG).

## Results

We obtained authentic ancient DNA from all individuals with between 0.9-14.1 % C>T nucleotide misincorporation on the 5’ end of sequenced DNA reads and 0.3-1.2 % G>A on the 3’ end, which is the expected pattern for single-stranded DNA libraries (see Table S1 and Figure S1). The mean length of mapped reads per sample ranged from 42.07 - 96.29 bp as is expected of short ancient DNA (Figure S1).

### Uniparental markers and sex determination

We were able to determine the genetic sex of all individuals and identified 10 males and 10 females. We recovered whole mitogenomes from 12 individuals and were able to call mtDNA haplogroups for 18 of the 20 individuals. Only two individuals had some mitochondrial contamination, at low levels (0.011–0.195; Table S1). All individuals carried Native American mtDNA haplogroups, with 12 individuals belonging to the C1b lineage, 3 to the D1 lineage and 3 to the D4h3a lineage. All 10 males carried the Native American Q1a2 Y chromosome lineage, with some belonging to more derived haplogroups. MtDNA and Y chromosome haplogroup assignments are reported in Table S1.

### Population structure in South America

The software ADMIXTURE was iterated in unsupervised mode 20 times for K values between 2 and 12 with K=7 showing the lowest cross validation values (Figure S2). At K = 7, three major South American ancestry clusters were identified, a broadly Andean component (yellow), a Patagonian component (green) and a Central Chilean component (pink) that is maximised in the ancient central Chilean individuals and modern Mapuche populations (Figure 2A). The Chono individuals display a larger proportion of the Patagonian component, and a smaller proportion of the Chilean component. Furthermore, previously published ancient Pampas individuals display a mixture of the Andes, Patagonia and Chile ancestry components. Three mid-Holocene Patagonian individuals and a 12,000-year-old individual from Central Chile are characterized by a larger proportion of the Andean component in comparison to more recent genomes available from the same regions. Considering the geographic locations and relative ages of these populations, it is likely that the ADMIXTURE results reflect an isolation-by-distance pattern rather than a real admixture between the ancestry components detected.

**Figure 2:**
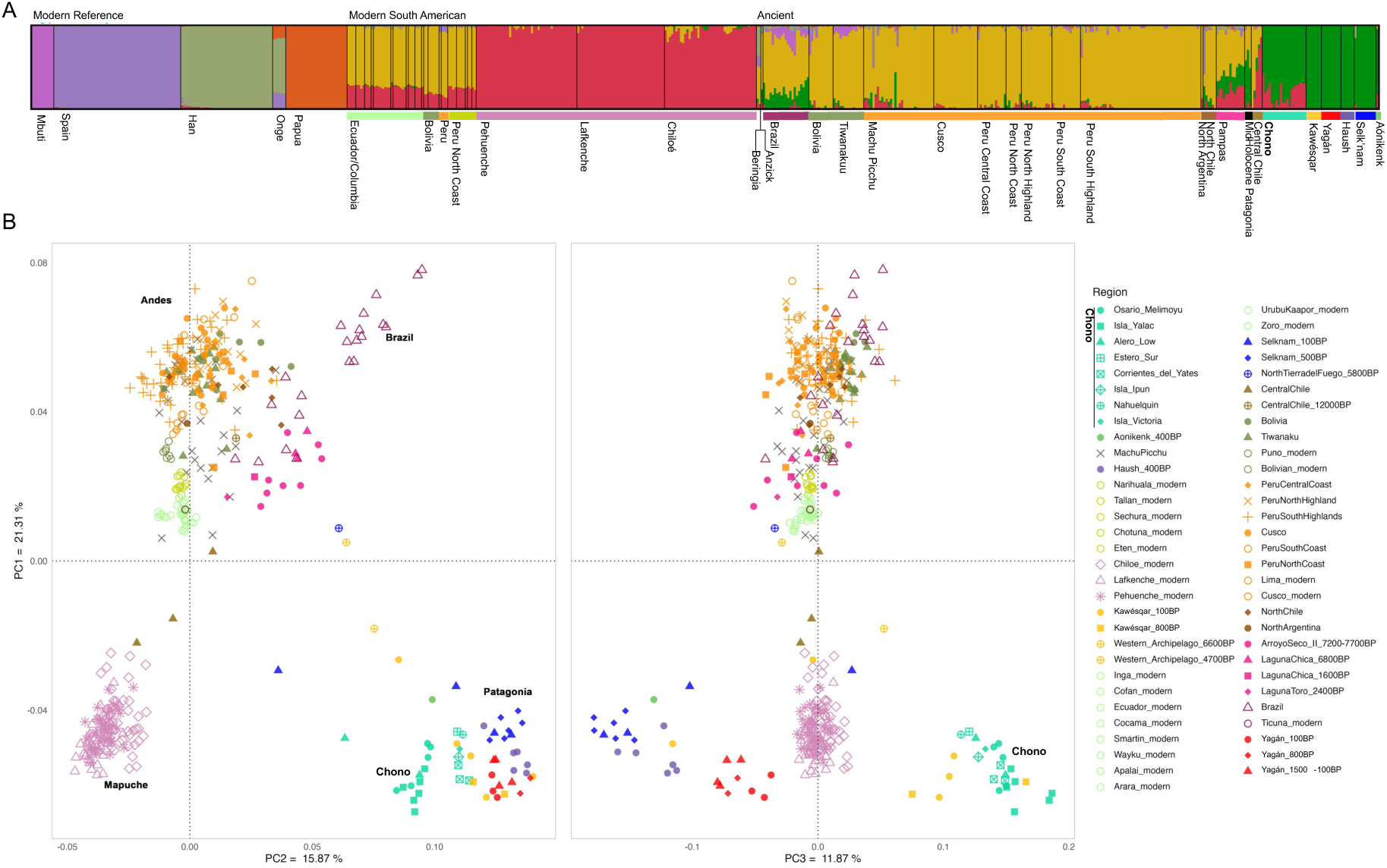
Population structure in South America. (A) Unsupervised ADMIXTURE result for K=7, indicated by lowest cross-validation value. Showing the mode of 10/10 iterations. Complete ADMIXTURE output in Figure S1. (B) PCA of modern and ancient South American individuals, showing the first three principal components. Calculated using EMU.

In the PCA, PC1 (21.31% of variation explained) represents a north–south cline in the South American continent, separating a cluster with Brazil and the Andes from a cluster with Chile and Patagonia (Figure 2B). Interestingly, PC2 (15.87% of variation explained) further separates the individuals from the Southern Cone according to their latitude, with one cluster containing ancient central Chile and modern Mapuche on the left, one cluster of Chono + Kawésqar in the middle, and another of far south Patagonia on the right. PC3 (11.87% of variation explained) separates the ancient Patagonian individuals into two clusters: one with the Chono + Kawésqar and one with the far south Patagonia. The MDS plot built with a transformation of the outgroup-*f*_3_ statistics (1-*f*_3_) (Figure S3) produced the same clustering, where now Dimension 2 corresponds to a north-south cline, and there is greater overlap between Patagonian groups, as MDS is by design more strongly influenced by the number of individuals per ancestry. In all plots, mid-Holocene Patagonian individuals fall toward the centre of the plot with more recent population clusters diverging away from them, corroborating that the observed population structure in Patagonia developed after this time.

We applied population outgroup-*f*_3_ statistics of the form *f*_3_ (Mbuti; Chono, X) to measure shared genetic drift between the Chono and other South American populations (Figure S4, Table S3). The populations experiencing most shared drift with the Chono are all the Patagonian groups. Within Patagonia, the level of shared genetic drift with the Chono correlates with the geographic distance along the Pacific coast from north to the southern tip of the continent and then back to north on the Atlantic coast (Figure 1A), such that Selk’nam and Aónikenk have the least shared drift, despite being geographically closer to the Chonos archipelago in a straight line across the land. The mid-Holocene Patagonian genomes (North Tierra del Fuego 5800 BP and Western Archipelago 4700 - 6600 BP) do not exhibit this high shared drift with the Chono, as other Patagonians later do, further suggesting the development of population structure in Patagonia after the mid-Holocene.

Clustering heatmaps plotted from pairwise individual *f*_3_ statistics (Figure S5-S6) reveal three main ancestry clusters within the SCSA. First, a “far-south” cluster, comprising Selk’nam, Yagán, Aónikenk and Haush; second, a “western archipelago” group containing the Kawésqar individuals clustered within the Chono individuals; and finally, a cluster of mid-Holocene genomes from the Pampas, central Chile and Patagonia (Figure S5). The “far-south” cluster is further divided into a “Selk’nam-like” cluster (north of Tierra del Fuego, also described as “foot nomads”) and “Yagán-like” cluster (south of Tierra del Fuego, also described as “sea nomads”), with Haush individuals distributed across both clusters (Figure S5). When modern Mapuche genomes are included in the clustering heatmap, they form one large cluster and ancient Central Chile individuals fall within it. The Pampas/mid-Holocene cluster is an outgroup to Patagonia, with the notable exception of one Chono individual CY5_1 clustering towards the Pampas group (Figure S6): the individual is described as a genetic outlier (see below).

### Genetic relatedness and diversity estimates within Chonos

We identified several close genetic relatives using BREADR ^55^, including two pairs from the archaeological site Isla Yalac 1. IYA1 (male) and IYA6 (female) have a first-degree relationship (Table S4-S5) and share the same haplotype belonging to mtDNA haplogroup C1b with (Table S1, S4-S5), suggesting they could be either full sibling brother and sister or mother and son. Both individuals are radiocarbon dated, and IYA6 (110-454 cal BP) has a date of approximately thirty years later than IYA1 (51-411 cal BP) could be in principle compatible with the mother-son hypothesis, although the range of radiocarbon dates is broader. Additionally, given the prevalence of the C1b haplogroup in this dataset, we cannot reject that the pair may be father and daughter who have coincidentally inherited the same C1b haplotype. A second-degree related pair was also identified, IYA3 and IYA5. Both individuals are male, with different mitochondrial haplogroups (C1b13 and C1b, respectively) and the same Y chromosome haplogroup (Q1a2a) (Table S1, S4-S5), suggesting they could be patrilineally related first cousins, half-brothers, uncle/nephew or grandfather/grandson. From the site Corrientes del Yates 5, we identified a second-degree relationship between CY5_3 and CY5_4, two females with different mitochondrial haplogroups (D4h3a5* and C1b, respectively). Similarly to the previous pair they could be first cousins or half-sisters not sharing their matriline, or granddaughter/paternal grandmother.

To further investigate the relationship between the two pairs from Isla Yalac 1, we tested for relatedness up to the 10th degree using BREADR ^55^, in all pairwise combinations of individuals from the sites of Isla Yalac 1, Corrientes del Yates 5 and Osario Melimoyu given they each had more than two individuals. Nearly all pairs tested returned a relationship between the 3rd and 6th degree (Table S5), which we do not interpret as a true signal of relatedness but as an effect of background relatedness, for example from small sustained effective population size bringing a higher chance of sharing relatives in the past.

We therefore analysed Runs of Homozygosity (ROH) using hapROH ^76^. All Patagonian populations tested had a high total length of ROH, when compared to ancient populations from further north in Bolivia, the Pampas (Laguna Chicha 1600BP) and North Argentina, (Figure 2A), in line with a small effective population size. Additionally, high total length of ROH is present in early Holocene genomes ArroyoSeco_II_7200-7700BP and CentralChile12000BP, indicative of small founding populations and possible serial founder effects. Throughout ancient Patagonia, there are high proportions of short ROH (4–8 cM) (dark blue, Figure 3A), which is suggestive of persistently small population sizes in the past ^76^. Strikingly, Chono individuals have an especially high amount of long 20–300 cM ROH (red, Figure 3A), suggesting more recent close-kin unions in the individual’s genealogies ^76^.

**Figure 3.**
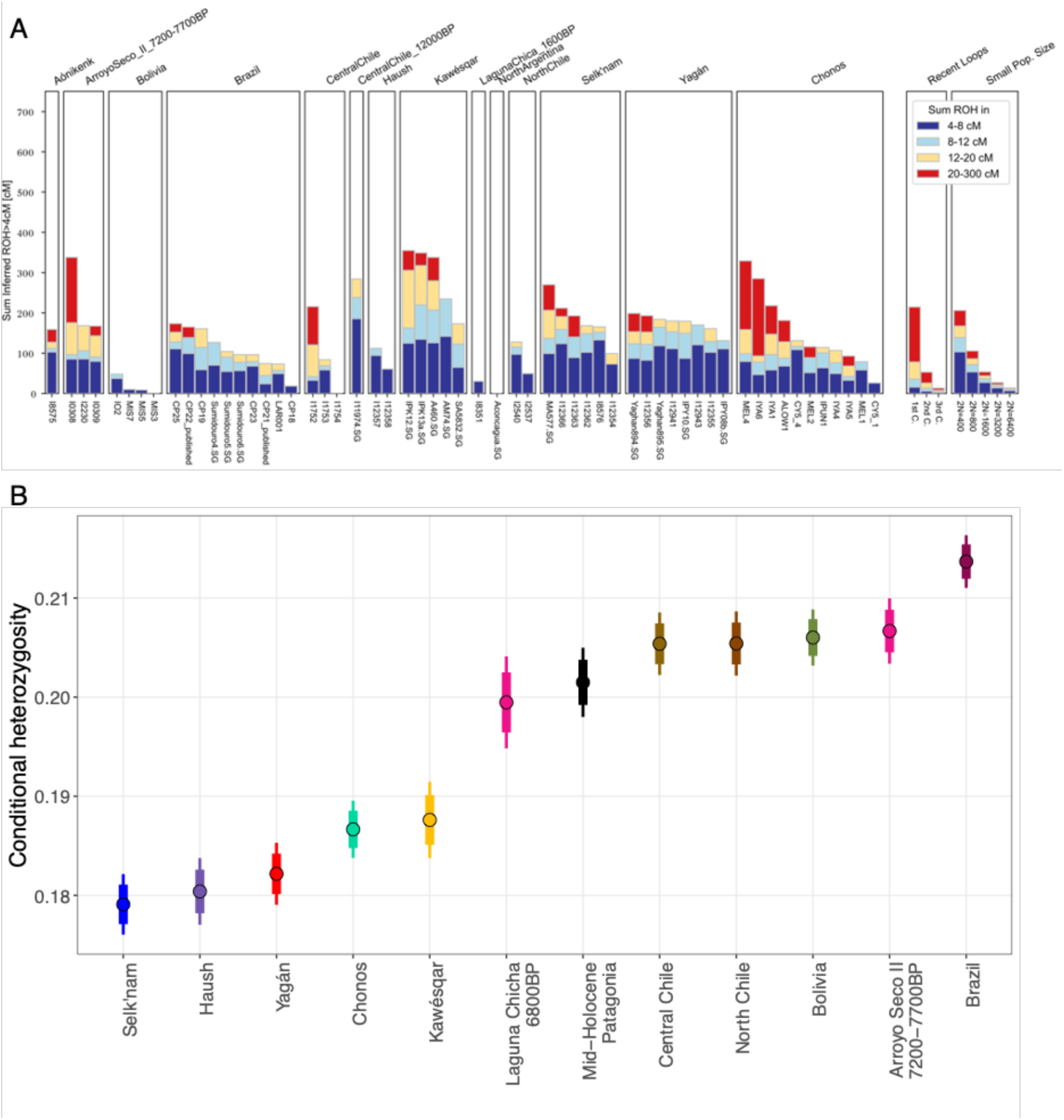
(A) ROH (runs of homozygosity) blocks identified in ancient Patagonian individuals with > 400,000 SNPs. The two right panels are estimated ROH for unions where the parents are full cousins of degree 1, 2, 3 and effective population sizes 200, 1000, 5000. (B) Conditional heterozygosity calculated within ancient populations where n > 1. Thick and narrow error bars represent ± 1.96 and 3.00 s.e., respectively.

To further understand the background relatedness in ancient Patagonian groups, conditional heterozygosity was estimated ^72^. The results corroborate previous reporting of low heterozygosity in Patagonian populations ^17^, and find that the Chono had low heterozygosity, similar to Kawésqar, but not as low as populations further south such as Yagán, Haush and Selk’nam (Figure 3B). This low conditional heterozygosity indicates persistently small population sizes and has likely developed since the mid-Holocene (8.2–4.2 kya), as Patagonian individuals from the mid-Holocene do not have this low level of conditional heterozygosity, and instead are similar to other ancient SCSA populations (Figure 3B). These results of large ROH and low heterozygosity in the southern regions of the southern cone contrasts reports of higher genetic diversity and smaller ROH lengths reported for living Mapuche populations in the regions immediately north of the Chonos archipelago (Arango-Isaza et al. 2023).

### Population structure in time and space in the Chonos archipelago

In order to examine structuring within the Chonos archipelago, we applied *f*_3_ statistics to test the shared genetic drift of each Chono individual to the modern Huilliche from Chiloé compared to the genetic shared drift with the ancient Kawésqar ancestry (Figure 4, Table S6). We can distinguish a north-south structure with “north-Chonos” from Isla Yalac 1 (IYA), Osario Melimoyu (MEL) and Alero Low (ALOW) having the least shared drift with Kawésqar, compared to individuals from sites further south in the Chonos (Figure 4). Though there is a clear relationship with latitude, there is also a correlation between the clustering pattern and time period. In fact, one limitation of this study is that each archaeological site represents a different time period, with the northern sites being more recent and southern sites being more ancient (Figure 1C). There is one exception: Nahuelquín 1 is a southern Chonos site with relatively recent dates of around 400 BP (Figure 1C). Unfortunately, the individual from Nahuelquín did not yield high quality genomic data (Table S1), and could not be included in further comparisons, making it difficult to explore the extent to which this structure within the Chonos archipelago is geographically or temporally derived.

**Figure 4:**
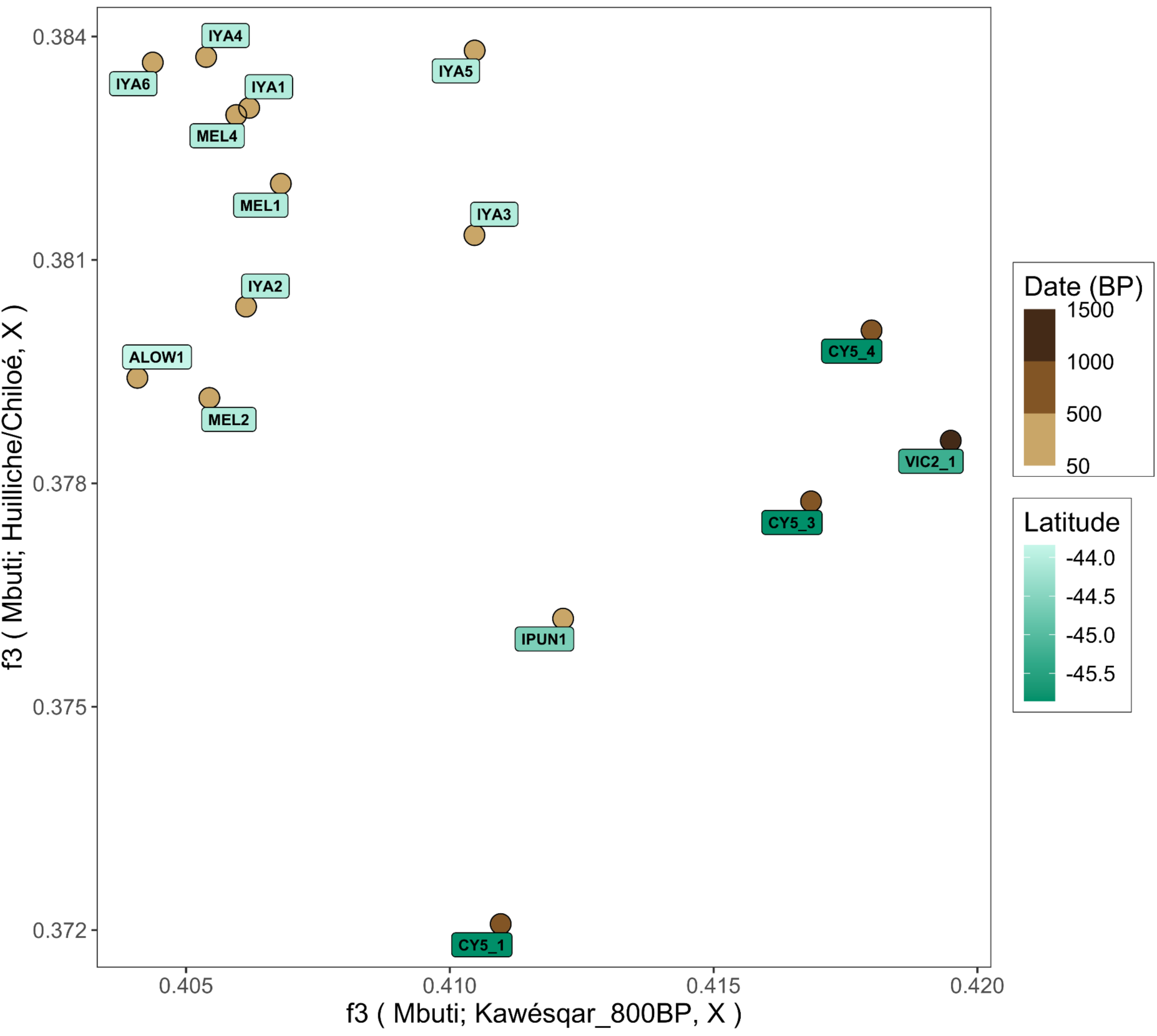
Comparisons of two *f*_3_-outgroup tests of the form *f*_3_(Mbuti; TestPopulation, X), where TestPopulation is ancient Kawésqar (x-axis) or modern Huilliche (y-axis), and X rotates each of the Chono individuals. Points are labelled by individuals ID, containing site specific abbreviation and label colour shows latitude. Point colour gradient shows date (BP) (right).

Using the north-south separation of Chono individuals emerging from Figure 4, we separated the Chono population into a Chonos North and a Chonos South subgroup (see Table S2 for population grouping details). We then tested the relative relationships to other South American populations with the statistic *f*_4_ (Mbuti, X; Chonos North, Chonos South) (Figure 5). Several populations have significantly more shared drift (|Z|>3) with Chonos South than Chonos North, including Kawésqar, Yagán - these two, with Chono are sometimes referred to as the “sea nomads” of Patagonia ^7^. Of the three Mapuche populations, only the Huilliche of Chiloé exhibit significantly more shared genetic drift (|Z|>3) with Chonos North rather than Chonos South, a pattern not seen in the other Mapuche groups, Lafkenche and Pehuenche.

**Figure 5:**
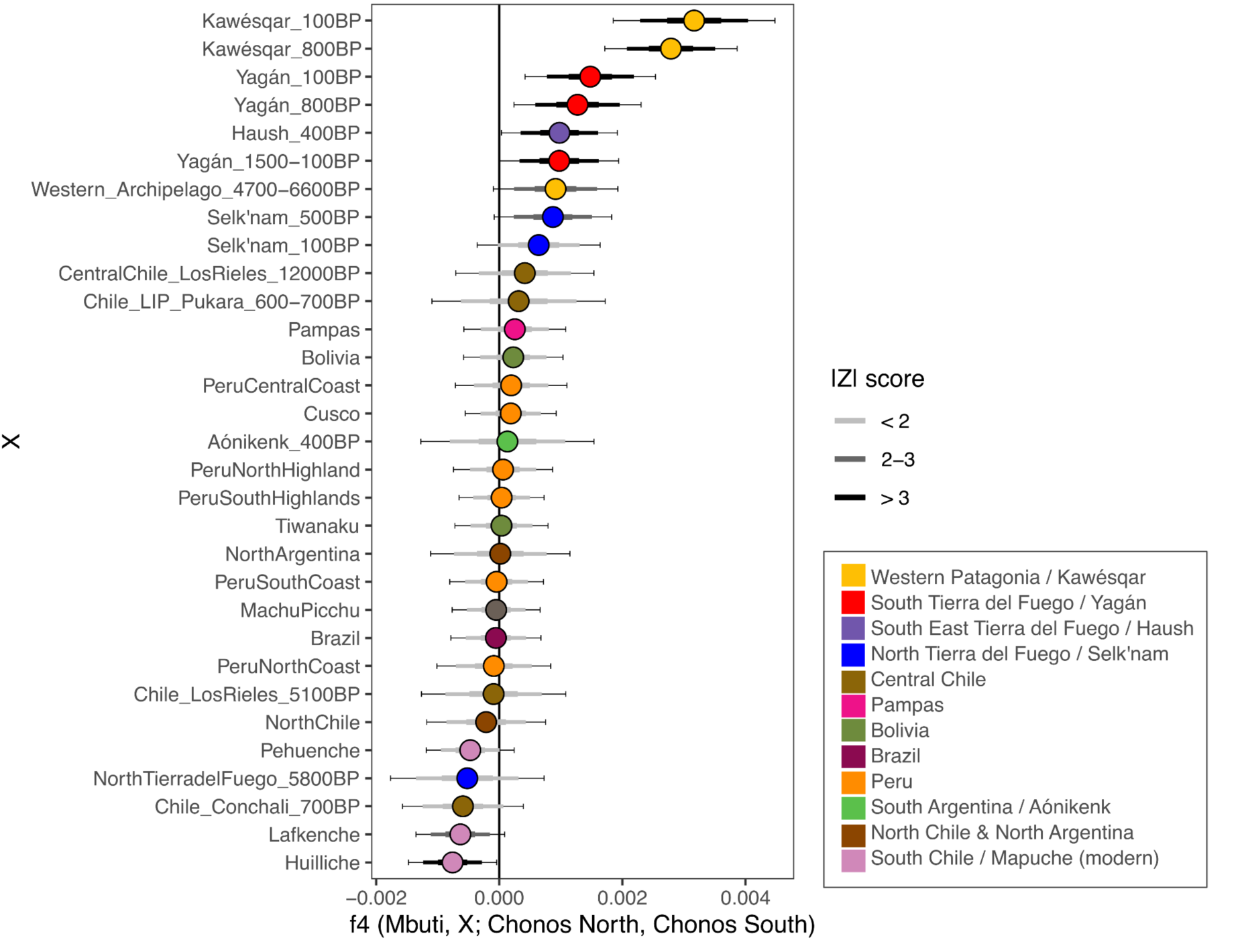
*f*_4_ statistics of the form *f*_4_ (Mbuti, X; Chonos North, Chonos South) examine differential relationship of Chonos North and Chonos South sub-groups with other South American populations in the X position. Point colour represents the broad geographical region. Error bars represent 3 s.e. And |Z| score is indicated by the colour of the bar.

### Connection between Chono and Mapuche

To further interrogate the relationship of Mapuche ancestries to the ancient Chonos ancestry we ran statistics of the form *f*_4_ (Mbuti, X; Huilliche, Lafkenche/Pehuenche) (Figure 6, Table S8). Chonos South does not exhibit significant shared drift with any of the three Mapuche groups. However, Chonos North has significantly more shared drift with Huilliche than Pehuenche (|Z| >3). Interestingly, Chile_Conchali_700BP and the ancient Pampas exhibit significantly higher shared drift with Lafkenche relative to Huilliche, but this is not found between Huilliche and Pehuenche. Pampas and Chile_Conchali_700BP are significantly closer to Lafkenche than Huilliche, but there is no significant difference between Huilliche and Pehuenche.

**Figure 6:**
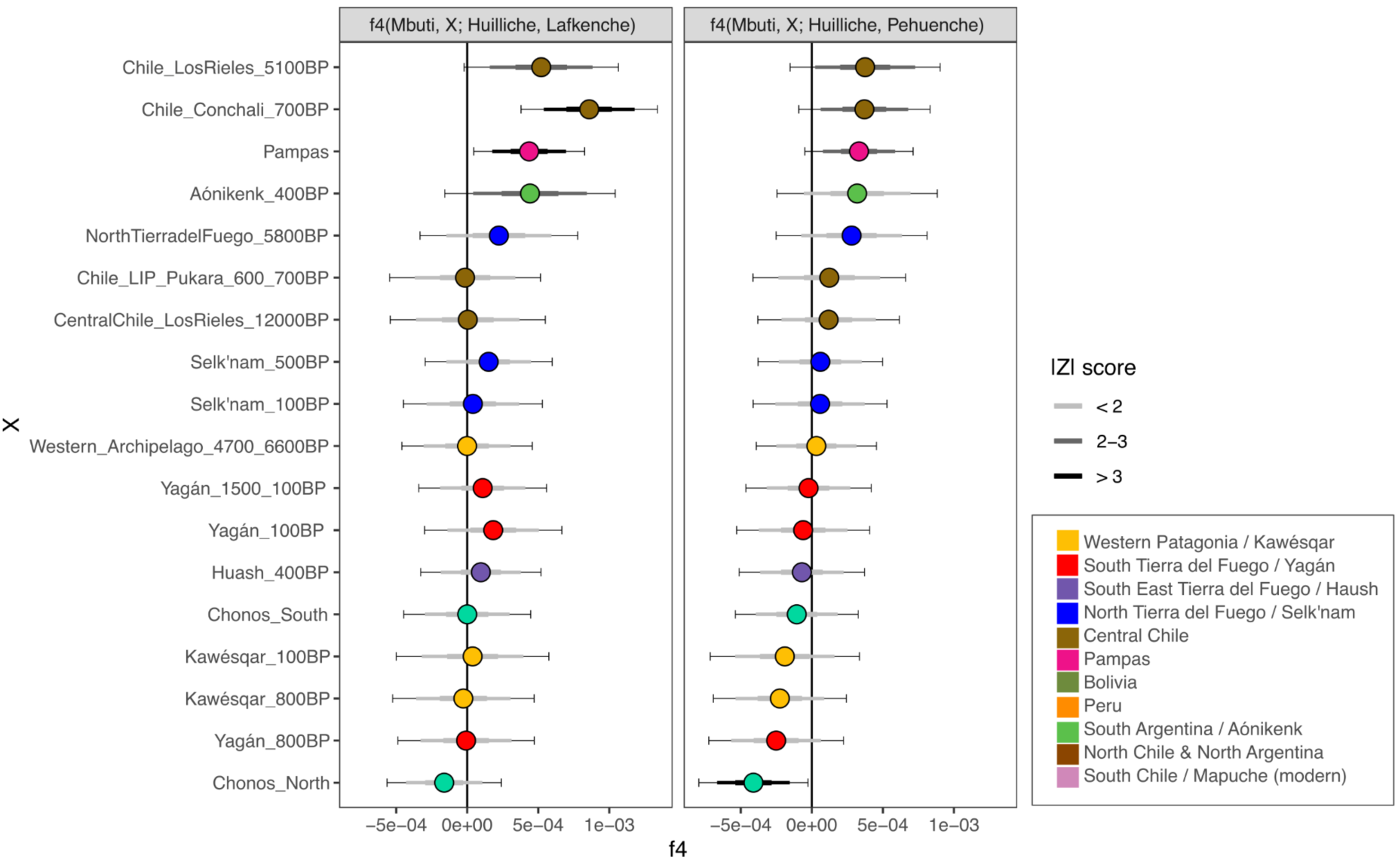
*f*_4_ statistics of the form *f*_4_ (Mbuti, X, Huilliche; Lafkenche/Pehuenche) to test the relationship of Mapuche groups to the Chono relative to their allele sharing with other South American populations in the X position. Point colour represents the broad geographical region. Error bars represent 3 s.e. And |Z| score is indicated by the colour of the bar.

Given indications of a connection between Huilliche and Chono, to test if this relationship was generally shared with all individuals or specifically in some individuals only, we ran *f*_4_ (Mbuti, Chono; A,B) (Figure S7) and *f*_4_ (Mbuti, Chonos North; A,B) (Figure S8), where A and B rotated pairwise between all Huilliche individuals. In both heat maps we can see some individuals have significantly more shared drift with the tested Chono population than their paired Huilliche individual, indicating overall that the relationship to the Chono affects some but not all members of the Huilliche population. Furthermore, to test the relative relationship of Huilliche/Chiloé individuals to the Chonos North cluster and to other Mapuche populations we performed *f*_4_ statistics of the form *f*_4_(Mbuti, Y; Chonos North, Lafkenche+Pehuenche), in this case combining Lafkenche and Pehuenche into one group, and rotating Y between modern Huilliche individuals (Figure S9, Table S9). We see that most Huilliche individuals have significantly more shared drift with the Lafkenche+Pehuenche grouping that with the Chonos North population (N=15), except one individual for whom *f*_4_ = 0, who therefore shares equal drift with both the Lafkenche+Pehuenche group and Chonos North. This result highlights also internal diversity inside the Huilliche individuals from Chiloé.

### The case of one outlier individual

Throughout several analyses the Chono individual CY5_1 shows a distinct profile from the rest of the Chono individuals, behaving as an outlier; it has relatively very few ROH (Figure 3A), is distant from the Chono cluster in the MDS plot (Figure S3), and rather clustered with the Pampas population in the large Southern Cone 1-*f*_3_ heatmap (Figure S6), yet in *f*_3_ statistics it has the least shared drift with the Pampas and Mapuche compared to other Chono individuals (Figure 4). Taken together, these results are inconclusive to define the ancestry of the individual: the individual in fact does not clearly belong to a different ancestry, so we cannot identify the individual is a migrant, but at the same time the individual seemed divergent from the Chono enough to suggest at least a partial relationship with a migrant.

Thus, we used qpAdm to test if the ancestry of the individual can be explained by a one or two-source models, exploring possible source ancestries with Kawésqar_800BP as the closest representative of later Western archipelago ancestry, MiddleHolocenePatagonia samples representing the basal ancestry of Patagonia, Chile_LIP_Pukara_600-700BP as an ancient central Chilean ancestry that is most similar to Mapuche populations and lastly, PeruCentralCoast samples as a representative of the Central Andean ancestry.

Results of the qpAdm analysis rejected all two-source models while all single-source models were plausible, with Kawésqar_800BP being associated with highest p-value (0.95, Table S10). This could be explained in several ways. First, CY5_1 is one of the oldest individuals analysed in this study, radiocarbon dated to 671-942 cal BP (Table S1) and therefore could harbour an older ancestral state of the Chono genetic lineages that differs from the later individuals. Second, CY5_1 may represent some residual undifferentiated southern Andean ancestry contribution to the archipelago, but that, given the presently available data, the individual cannot be clearly connected to a specific contributing population or region. Thirdly, the analysis setup simply has minimal power to distinguish between ancient South American ancestries which are characterized by a certain degree of homogeneity across the continent - at least with the 1.2 million SNP set used here. In this light, whole genome data might provide further resolution to distinguish different ancestry contributions to a finer scale.

## Discussion

### Maternal ancestries of the Chonos archipelago

The mtDNA haplotypes detected in the newly sequenced Chono ancient individuals belong to three South American lineages, C1, D1, and D4h3a (Table S1). The same haplogroups were retrieved in a recent study of mtDNA diversity of the Chono population ^18^. We do not find haplogroups A2 or B2, consistent with previous studies that found these as nearly absent in populations of the extreme south ^85–87^. The most common haplogroup in our sample is C1b haplotypes (66 %), which is also the most common in the ancient Chono individuals described by Moraga and colleagues ^18^. We also identify three individuals within the Pan-American minor founding lineage D4h3a, which is found at highest frequencies in living populations from Patagonia and Tierra del Fuego ^88,89^. Finally, we identify three individuals carrying the same D1 haplotype, with one of two diagnostic SNPs for the D1g haplogroup ^90,91^, which is found nearly exclusively in the SCSA, most frequently in Argentinian and Chilean Patagonia, as well as in the Argentinean Pampas ^89,90,92^. This raises the potential that they are a haplotype basal to D1g, or D1g haplotypes with a back mutation.

### Population structure of ancient Patagonia

Our results replicate the previously reported genetic structure of SCSA from the late Holocene SCSA ^12,17,25^, which can be described with an isolation-by-distance ecological model, with geographically correlated clines as shown in ADMIXTURE, PCA (Figure 2) and MDS (Figure S3). In particular, we corroborate the observation of an ancestry cline whereby individuals are ancestrally most similar to those geographically closer to them along the Patagonian coast (Atlantic or Pacific) within the last two millennia ^17^ and expand this pattern up to the Chonos archipelago region, which is here described as the northernmost boundary of the ancestry characteristic of the southern region of the southern cone. Within this Patagonian ancestry, the Chono peoples are genetically most similar to ancient Kawésqar from the immediate south (Figures 2, 5, S3-S6).

The Chonos archipelago marks the northern boundary of the extension of the southern Patagonian ancestry. This ancestry is related, but slightly distinct, to the one characterizing Mapuche populations. The two regions also correspond to slightly distinct demographic histories, as Mapuche populations exhibit genetic signatures of a larger population size in comparison to Chono and other Patagonian groups, with the latter characterized by a north-south gradient of increased geographic isolation and genetic drift. In Patagonia, the Chono population has a similarly low heterozygosity estimate as Kawésqar ^93^, but not as low as the far south populations of Selk’nam, Haush and Yagán, suggesting a correlation between relative geographic isolation and small population sizes with limited gene flow that may result in low estimates of conditional heterozygosity. This genetic connection between ancient Chono and other sea nomads from southwest Patagonia suggests a similar demographic history for seafaring cultures of the Pacific coast. The smaller population size could be correlated to the harsh environmental conditions of the southern regions, which might provide a smaller carrying capacity. The reduction in population size could also account for the drastic effect of post-European contact decimating the Indigenous groups ^9,10,14,33^.

Finally, we report fine-scale resolution diversity patterns, which separate two clusters within the archipelago: a southern one, comprising more ancient individuals, and a northern one, comprising more recently dated individuals (Figure 4). The southern cluster has a greater shared drift with neighbouring Kawésqar individuals from the south, while the northern cluster shows excess shared drift with present-day Huilliche from the neighbouring Chiloé island in the north.

### Connections between Chono and Mapuche populations

The preferential sharing between aDNA profiles from the northern islands of the Chonos archipelago and current inhabitants of the island of Chiloé can be contextualised with historical and cultural evidence. Archaeological findings and ancient toponyms in Chiloé testify to a northernmost Chono presence in the island. Chono culture contributed a possible ancient substrate in shaping the identity of the island together with the Mapuche/Huilliche culture until today, an element of which part of the population is aware of, even if the circumstances of this encounter of peoples and cultures are not fully resolved. Linguistic and ethnographic surveys collected by from Jesuit missionaries in 1612 to an English ship captain in 1875 reported that in the island Canoe nomads had adopted a few words from Araucanian (Mapuche) languages ^9^ and other cultural elements possibly from Mapuche groups like sporadic gardening (e.g. potatoes) and herding, the polished stone axe and the plank boat (dalca). There is agreement that the Chono language was certainly distinct from the one spoken by Mapuche (Mapudungun) and Tehuelche (Chonan language family, distinct from Chono), and more probably than not also from Kawésqar ^94^.

In a previous IBD sharing analysis with modern DNA, the Huilliche/Chiloé population displayed a high amount of short and long IBD blocks shared with Kawésqar, pointing at a specific connection with the southern population that was not found in the Lafkenche and Pehuenche ^25^. Additionally, there has been identified a sphere of Huilliche influence reaching southern Chile, and with sea-faring peoples ^16,95^. These findings can be paired with our *f*_4_ statistics supporting a connection between the Chonos North and Huilliche/Chiloé that does not extend to Lafkenche and Pehuenche (Figure 5 & 6). Note that we cannot refute the possibility that our comparison between the two populations may be impacted by ascertainment bias in the SNP array platform used to genotype the modern DNA dataset, which has a lesser number of variants in comparison to the capture employed in our aDNA study, or genetic drift occurring in the past few centuries since colonisation.

In conclusion, we confirm the existence of a Chono ancestry cluster, which is most similar to the genetic ancestry of other maritime groups, within the broader context of Patagonian ancestries. The Chono ancestry cluster exhibits a north-south gradient, and we identify genetic links between the northernmost Chono individuals and the Huilliche individuals from the island of Chiloé.

## Author Contributions

Conceptualisation - RD, BL, LFS, OR, CB, EAI, Lab - KK, RD, LFS

Data Processing - RD, LFS

Data Analysis - RD, LFS, XRR

Manuscript writing - RD, BL, LFS, CB, XRR

Manuscript editing - All authors

Funding - LFS, BL, RT

Sample Provision - OR

## Acknowledgements

The greatest thanks go to the human beings, both present and past, whose genomes form the basis of this research, and all those that made their participation possible. Laboratory work at The University of Adelaide was conducted with technical support from Vilma Pérez and Corrine Mensforth and computational analyses were conducted using supercomputing resources provided by the Phoenix HPC service at the University of Adelaide with support from Fabien Voisin. We thank Raymond Tobler who contributed funding for laboratory work. We thank Víctor Moreno-Mayar and Rosa Fregel for insightful comments on an earlier version of this manuscript that was presented as a chapter in Roberta Davidson’s PhD thesis. This research was supported by The University of Adelaide’s funding to RD; the Agencia Nacional de Investigación y Desarrollo, Chile, (ANID-FONDECYT 1210045, 1260247, ANID-Regional R20F0002, ANID-Basal FB210018) to OR; the National Science Foundation (NSF - 1515138) to LF-S; Australian Research Council Future Fellowship (FT170100448) to BL; the NCCR Evolving Language, Swiss National Science Foundation Agreement 51NF40_180888 to CB.

## Supplementary Information

Supplementary Document 1 - Supplementary Figures

Supplementary Document 2 - Report in Spanish

Supplementary Document 3 - Supplementary Tables

