## Supplementary Document 1 for "Palaeogenomics reveals 1,500 years of population history of the peoples of the Chonos Archipelago, Chile"

### Ancient DNA authentication

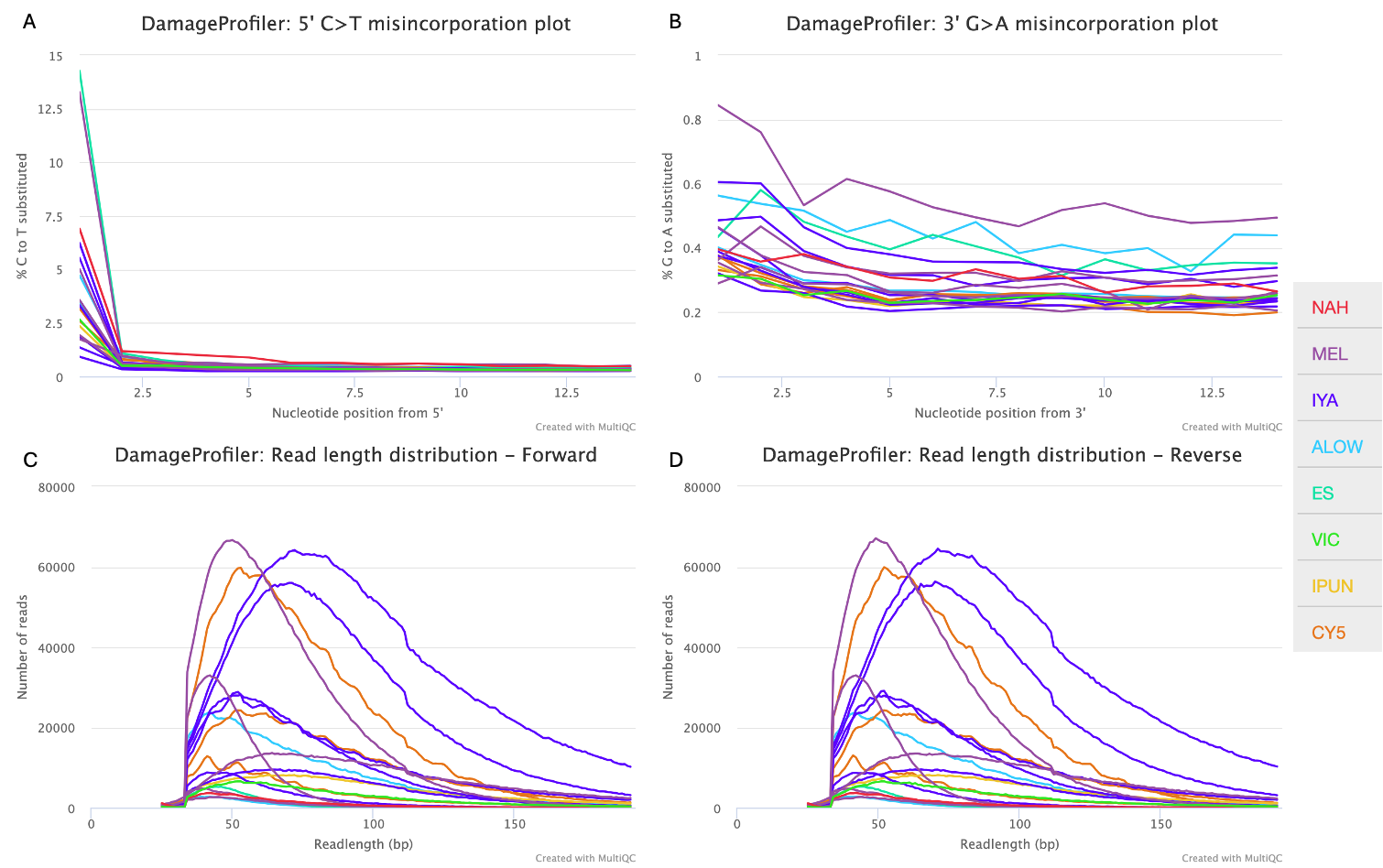

**Figure S1:** Ancient DNA damage in ancient Chonos libraries by DamageProfiler (ref). (A) 5’ C>T misincorporation by position, (B) 3’ G>A misincorporations by position, (C) read length distribution of the forward read and (D) read length distribution of the reverse read. Data used to generate plots is reported in Table SX. Samples are colour coded by site.

### Population Structure

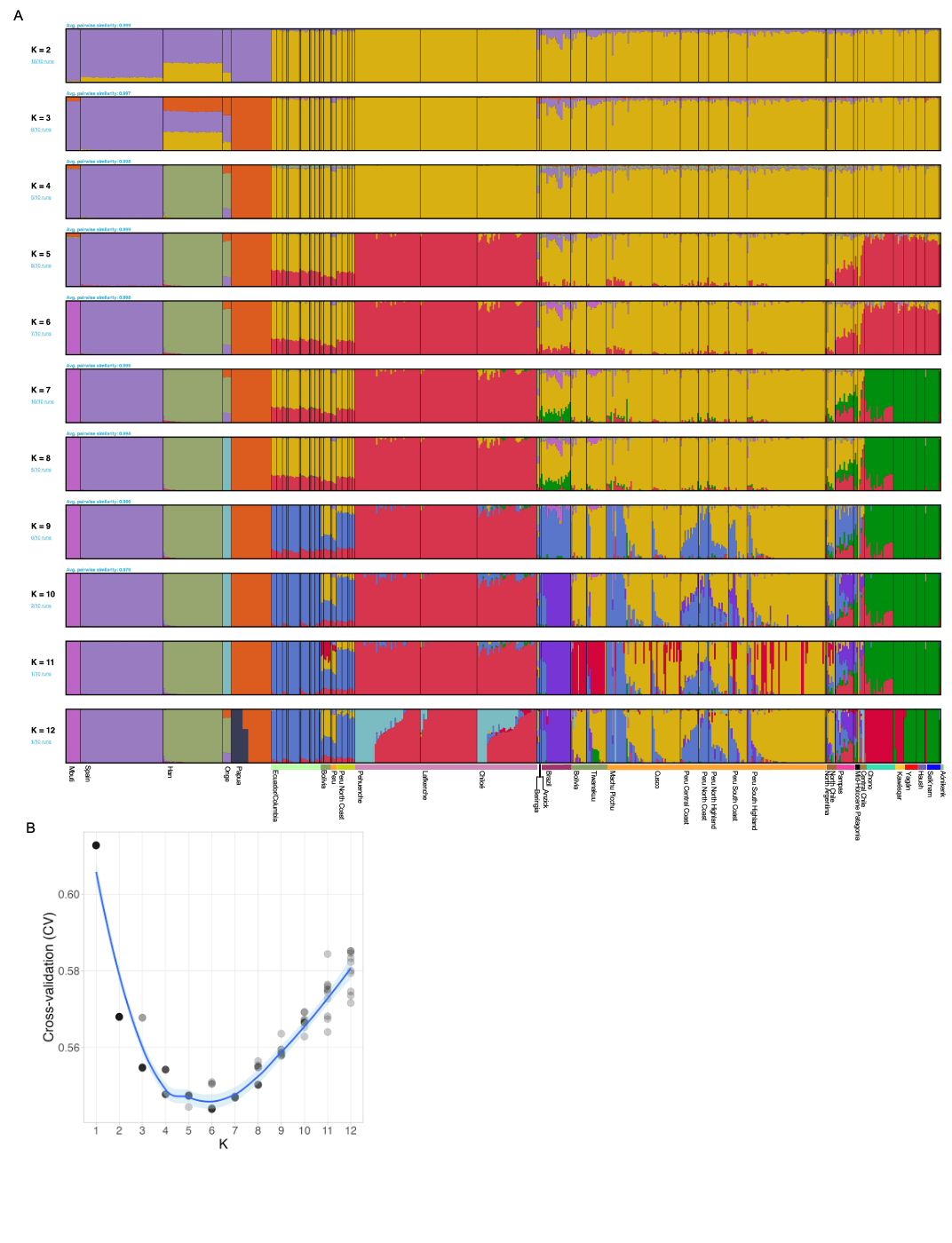

**Figure S2:** (A) ADMIXTURE results for K = 2 to K = 12, 10 iterations. (B) Cross-validation plot.

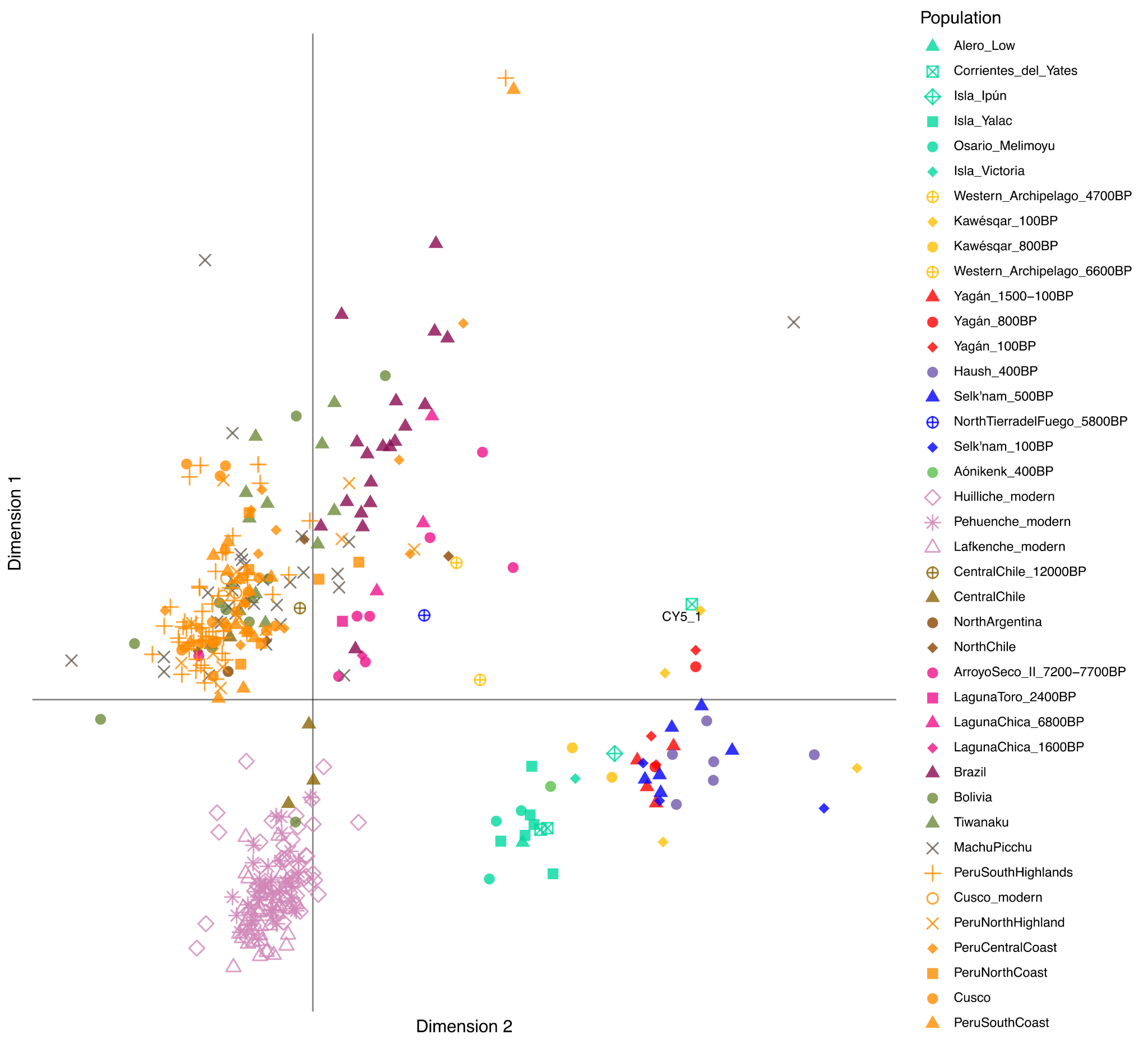

**Figure S3:** Multidimensional scaling (MDS) plot of the matrix of statistics 1-*f*_3_(Mbuti; Ind1, Ind2), where Ind1 and Ind2 are modern and ancient genomes from South America. Individual CY5_1 is described as a genetic outlier of the Chono group.

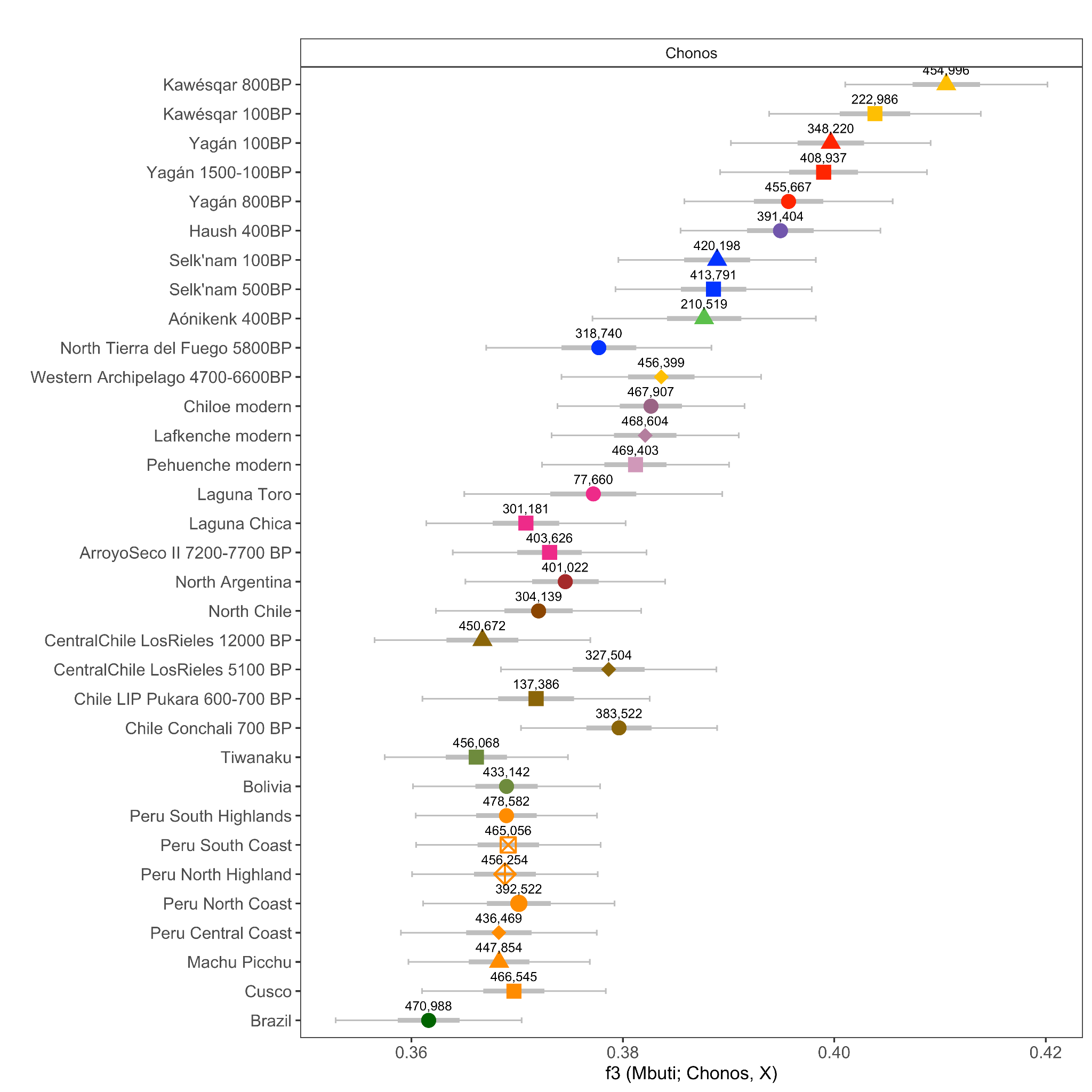

**Figure S4:** Outgroup *f*_3_ statistics of the form *f*_3_ (Mbuti; Chonos, X), where X rotates each South American population used in the study. The number of SNPs used in the statistic is annotated above the point. Thick and thin error bars show 1 and 3 s.e. respectively.

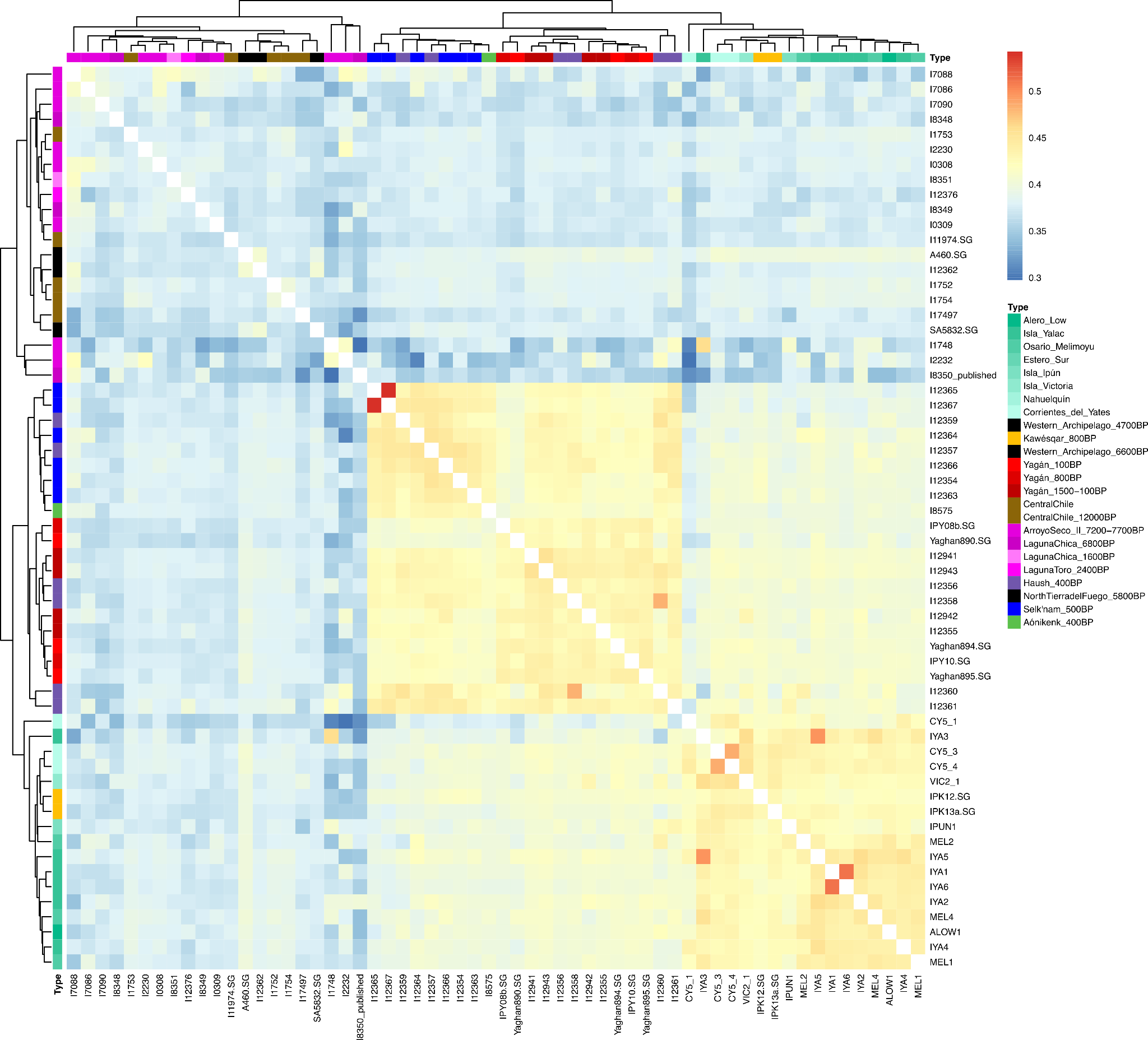

**Figure S5:** Heatmap of outgroup *f*_3_ pairwise comparisons of the form *f*_3_ (Mbuti; Ind1, Ind2) for ancient Patagonian individuals, annotated by population group. Individuals with < 1000 SNPs and non-Chono individuals dated post-contact were removed. Clustering performed in R using the pheatmap package to draw a cladogram (Kolde, 2019).

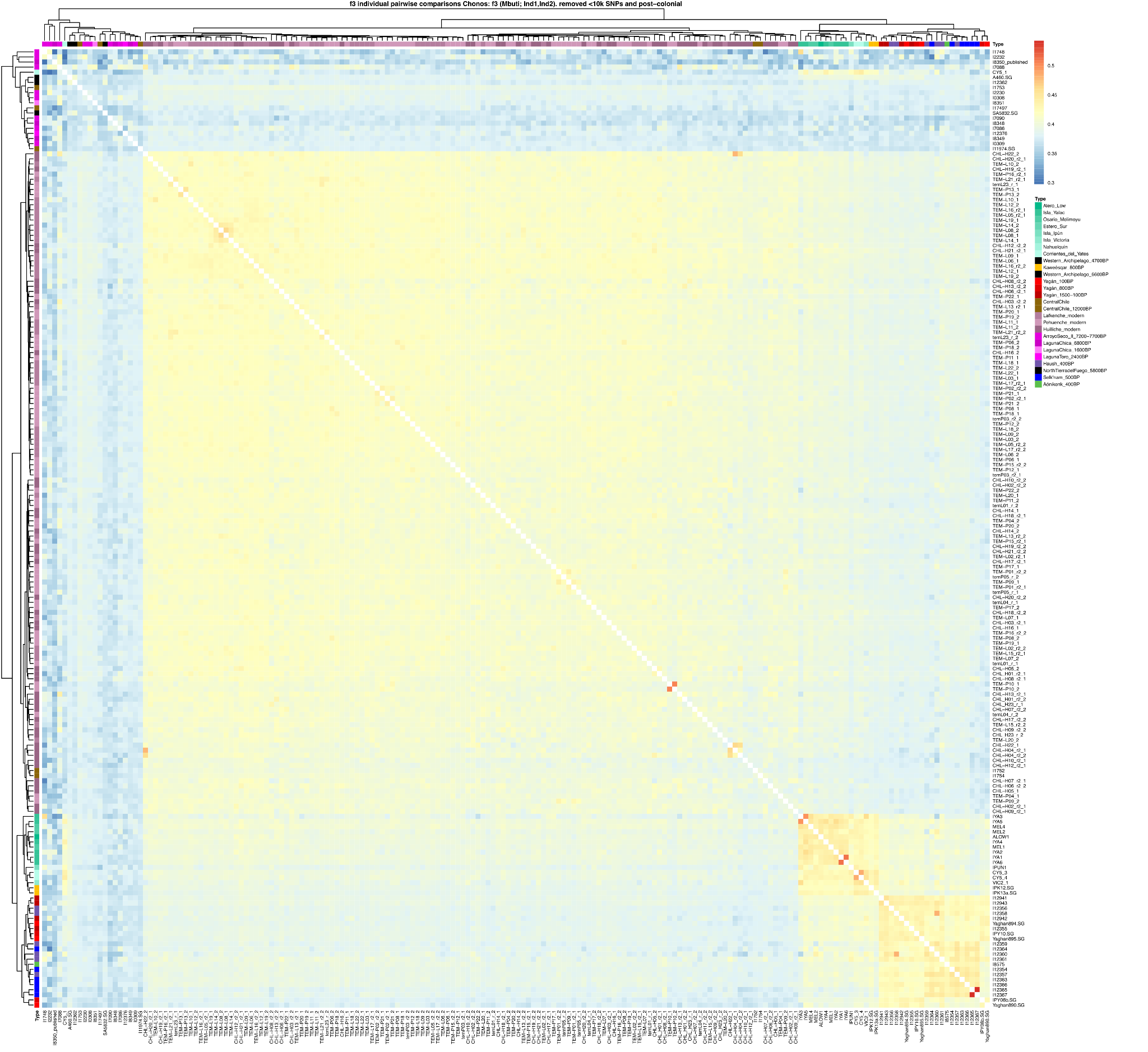

**Figure S6:** Heatmap of outgroup *f*_3_ pairwise comparisons of the form *f*_3_ (Mbuti; Ind1, Ind2) for Southern Cone individuals, including pre-colonial ancient individuals and modern mapuche individuals, annotated by population group. Ancient individuals dated post-colonisation or with < 1000 SNPs were removed. Clustering in R using the pheatmap package to draw a cladogram (Kolde, 2019).

### Huilliche-Chonos connection

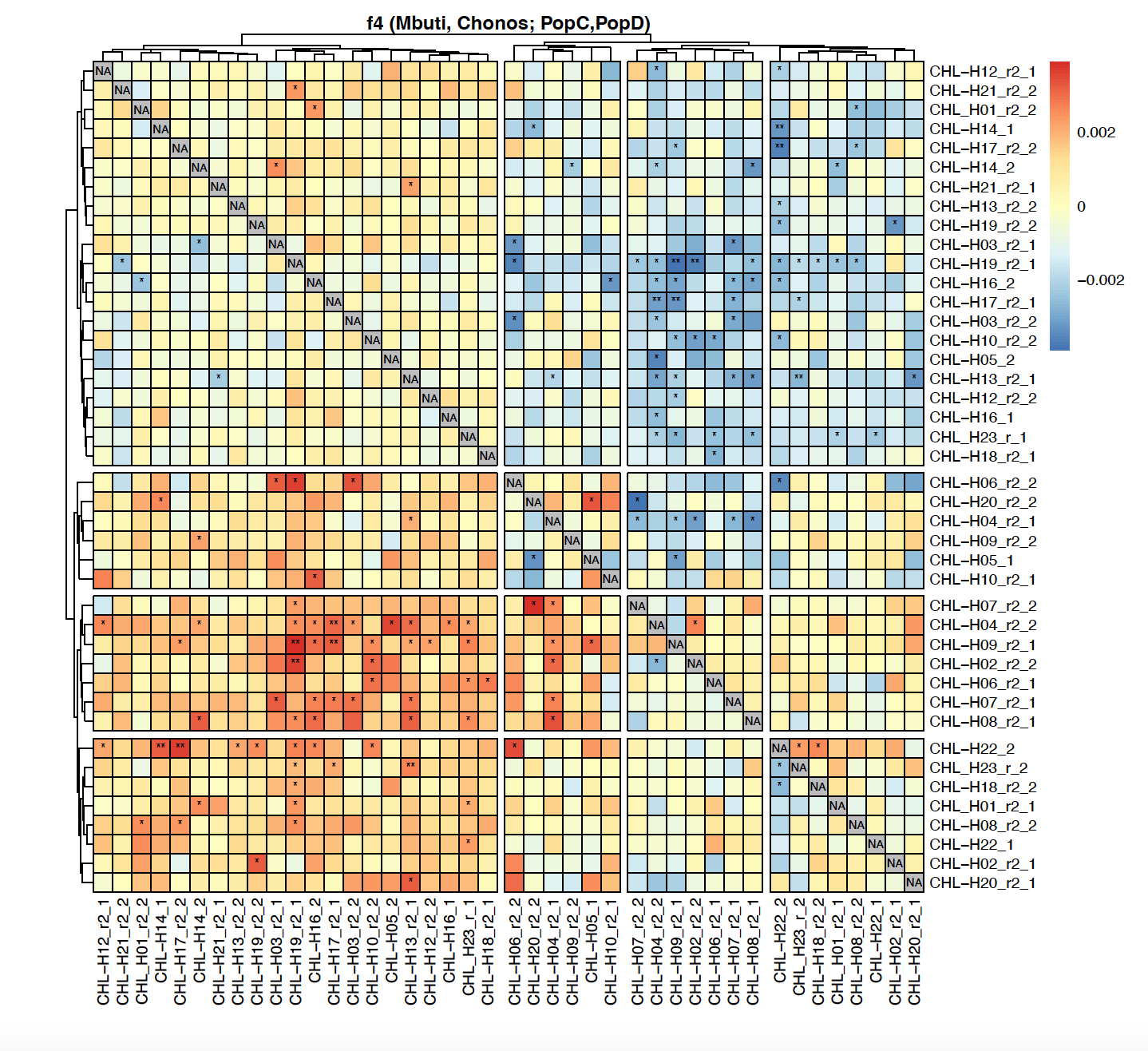

**Figure S7:** Heatmap of *f*_4_ statistics of the form *f*_4_ (Mbuti, Chonos; Ind1, Ind2) to investigate the relationship of Huilliche/Chiloé individuals to the whole Chonos population, where Ind1 and Ind2 rotate individuals from Huilliche/Chiloe population and Chonos is all Chonos individuals.

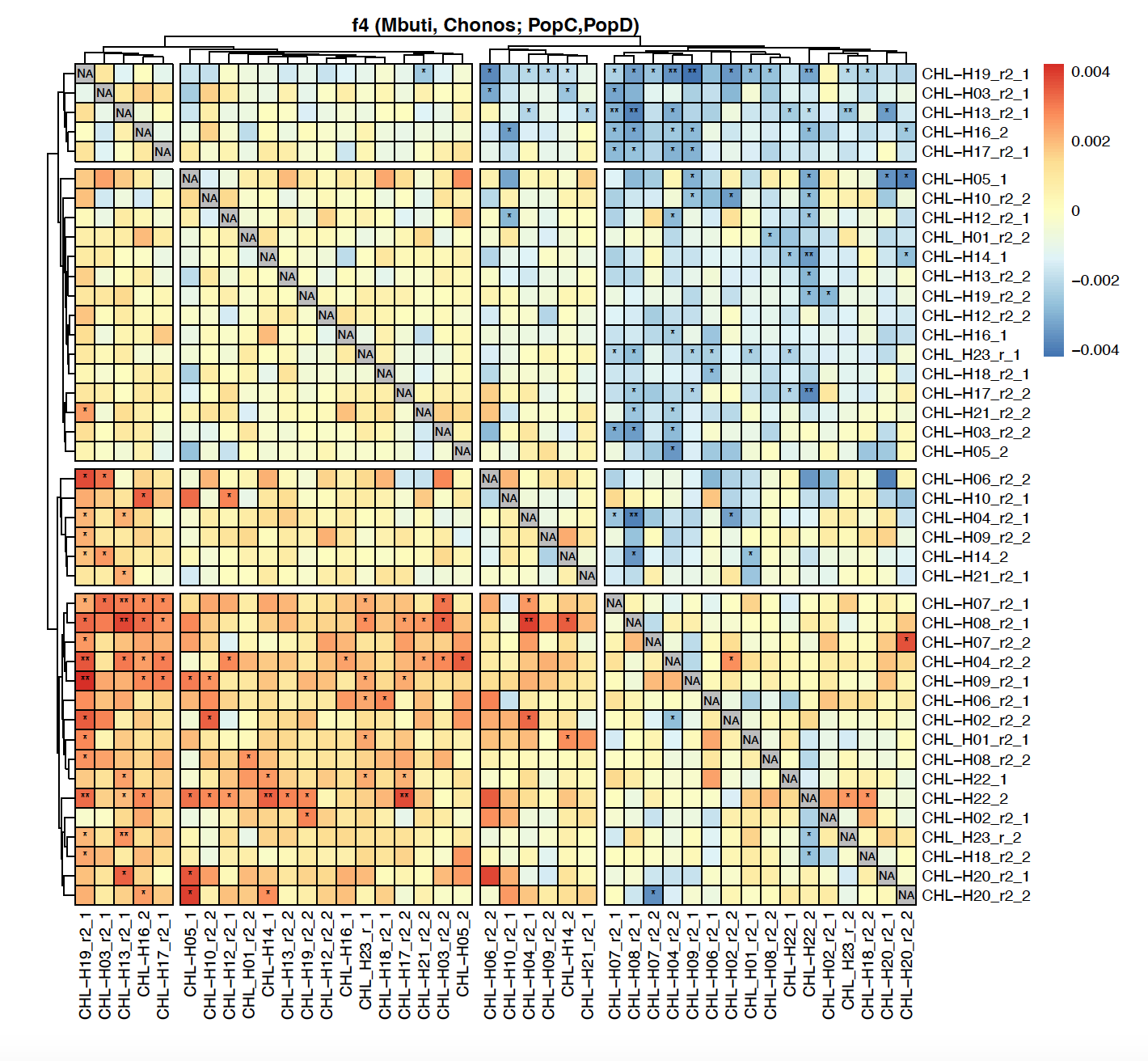

**Figure S8:** Heatmap of *f*_4_ statistics of the form *f*_4_ (Mbuti, Chonos_North; Ind1, Ind2) to investigate the relationship of Huilliche/Chiloé individuals to the whole Chonos population, where Ind1 and Ind2 rotate individuals from Huilliche/Chiloe population and Chonos_North is all individuals assigned to the Chonos North group.

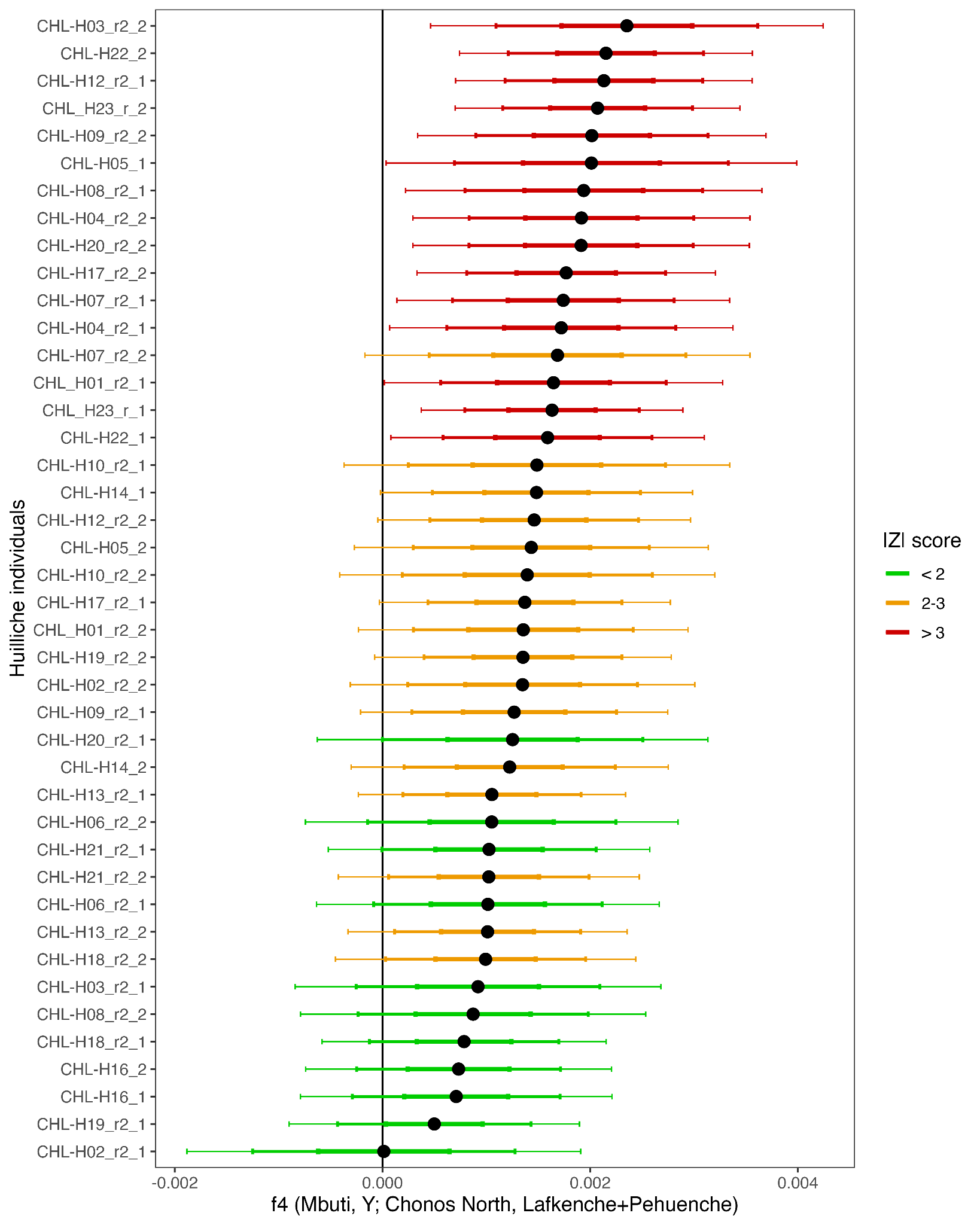

**Figure S9:** Statistics of the form *f*_4_ (Mbuti, Y; Chonos North, Lafkenche+Pehuenche) where Y rotates between modern Huilliche individuals
